# The most developmentally truncated fishes show extensive *Hox* gene loss and miniaturized genomes

**DOI:** 10.1101/160168

**Authors:** Martin Malmstrøm, Ralf Britz, Michael Matschiner, Ole K. Tørresen, Renny K. Hadiaty, Norsham Yaakob, Heok H. Tan, Kjetill S. Jakobsen, Walter Salzburger, Lukas Rüber

**Author notes:** Corresponding authors Ma.M. and L.R.

## Abstract

*Hox* genes play a fundamental role in regulating the embryonic development of all animals. Manipulation of these transcription factors in model organisms has unraveled key aspects of evolution, like the transition from fin to limb. However, by virtue of their fundamental role and pleiotropic effects, simultaneous knockouts of several of these genes pose significant challenges. Here, we report on evolutionary simplification in two species of the dwarf minnow genus *Paedocypris* using whole genome sequencing. The two species feature unprecedented *Hox* gene loss and genome reduction in association with their massive developmental truncation. We also show how other genes involved in the development of musculature, nervous system, and skeleton have been lost in *Paedocypris,* mirroring its highly progenetic phenotype. Further, we identify two mechanisms responsible for genome streamlining: severe intron shortening and reduced repeat content. As a naturally simplified system closely related to zebrafish, *Paedocypris* provides novel insights into vertebrate development.

## Introduction

The developmental mechanisms that determine how genotypes translate into phenotypes and how selection acts on genotypes to shape morphological differences are central to our understanding of the diversity of living organisms^1,2^. Model organisms have been instrumental in our quest to decipher how genetic differences are associated with morphological and physiological disparity^3^, and improved technologies for genetic modifications will aid in further elucidating genotype-phenotype interrelations^4^. One interesting example is the recent work by Nakamura *et al.* (2016), which utilized CRISPR-Cas9 to knock out three *Hox13* copies (*Aa*, *Ab*, and *D*) in zebrafish (*Danio rerio*) to investigate the role of these genes in the transition from fins to limbs. Although experiments like these are now feasible, the pleiotropic effects of genes involved in fundamental developmental processes, such as those of the large *Hox* gene family, present challenges and limitations to the phenotypic variation that can easily be induced in model organisms through genetic engineering. Examining naturally occurring extreme phenotypes in close relatives of model organisms thus provides a novel source of phenotypic and genotypic variation that is becoming increasingly important in improving our understanding of the molecular basis of evolutionary changes^5^.

Discovered around 200 years ago^6^, the zebrafish has been used as a molecular model organism since the 1980s and is currently one of the most important model systems for studying vertebrate development, genome evolution, toxicology, physiology, behavior, and disease^7,8^. Comparative efforts have so far focused on closely related *Danio* species^9,10^, but recent studies have revealed that the diversity of zebrafish mutants is surpassed by the range of phenotypic variation among several of its related species in the wild^3,11,12^. Some of these other members of Cyprinidae (e.g *Paedocypris, Sundadanio*, and *Danionella*) are characterized by developmental truncation^13–16^ and morphological novelties^16–18^ and thus offer additional, underappreciated potential for comparative studies and promise fundamental and transformative advances in our understanding of the molecular underpinnings of evolutionary change leading to novelty and adaptation at a genetic and phenotypic level^3^.

The Series Otophysi, which includes Cyprinidae, represents one of the earliest diverging teleostean lineages, and the phylogeny of this lineage remains controversial^13,19-23^. One particularly difficult taxon to place confidently is the recently discovered miniaturized dwarf minnow genus *Paedocypris*^13,15,21,22,24^ found in the highly acidic blackwater of endangered peat swamp forests in Southeast Asia. This genus of tiny vertebrates includes the world’s smallest fish, maturing at ~8 mm^25^. *Paedocypris* exhibits an extreme case of organism-wide progenesis or developmental truncation, resulting in an anatomical adult condition closely resembling that of a 7.5 mm zebrafish larva, with over 40 bones not developed^15^. To investigate the genomic signatures of developmental truncation, we sequenced and compared the genomes of two representatives of the genus *Paedocypris*; *P. carbunculus* and *P. micromegethes*. The genome signatures of these two species allow the distinction of genus-specific from species-specific genomic changes and thus enable identification of features associated with the extreme phenotype of *Paedocypris*. By comparing their genomes with those of other teleosts, including the closely related zebrafish, we identify *Paedocypris*-specific genomic signatures of developmental truncation in the form of loss of various key developmental genes, mirroring their progenetic phenotype. We further demonstrate how the *Paedocypris* genome size has been reduced through shorter genes due to significantly shorter introns compared to zebrafish, while exon lengths and gene numbers have remained relatively unchanged. We find that the *Paedocypris* genomes are comparable in size and gene structure (i.e short introns and compact genomes) to those of the two pufferfish model organisms; *Takifugu rubripes* and *Dichotomyctere nigroviridis*, which have the smallest vertebrate genomes known^26^. Additionally, we show that the accumulation of transposable elements (TEs), especially DNA transposons, is very low, following the diversification of the genus *Paedocypris*, and propose a potential mechanism for enhanced transposon-silencing activity through duplication of the PIWI-like (*PIWIl1*) gene in this lineage. Highly progenetic fish species like *Paedocypris* will be important resources for future studies on vertebrate development, presenting a novel opportunity to investigate phenotype–genotype relationships in early vertebrate development and the genetic mechanisms of developmental truncation.

## Results

### Loss of *Hox-* and other developmental genes

*Hox* genes encode transcription factors essential in body patterning along the anterior-posterior body axis during early development of all animals^27^. Although a large number of genes is involved in developmental processes, *Hox* genes are especially interesting as they remain organized in conserved clusters and are ordered along the chromosomes according to where and when they are activated^28^.

As all known *Paedocypris* species have a progenetic phenotype, we investigated whether this condition is reflected in their *Hox* gene repertoire. We compared 70 *Hox* gene transcripts, encoded by the 49 *Hox* genes in zebrafish, to detect syntenic *Hox* cluster regions in the two *Paedocypris* genomes (Methods Supplementary Note and Supplementary Table 1). For *Hox* clusters *Aa*, *Ab*, and *Bb*, we recover all genes on one to three contiguous sequences in synteny with *Danio rerio* (Figure 1a). Two scaffolds in both *Paedocypris* species cover the *HoxBa* cluster; however, functional copies of *hoxB10a* and *hoxB7a* cannot be identified in this region. In total, 10 of the zebrafish *Hox* genes appear to be absent in *Paedocypris*, and their predicted positions are illustrated as red lines in Figure 1a. While the remaining *Hox* clusters are recovered as more fragmented, most of the genetic regions are covered by contiguous sequences containing intact copies of other *Hox* genes in at least one of the two *Paedocypris* species. None of the 10 *Hox* genes could be detected in other parts of the genomes by sequence similarity searches, strengthening the hypothesis that these genes are indeed lost in this lineage. Although extensive *Hox* gene loss has been reported for other cyprinids, these are secondary losses, following genome duplication events^29^. Figure 1b shows the positions of both the missing and the present *Hox* genes in *Paedocypris* compared to currently available otophysan genomes, illustrating the most plausible reconstruction of *Hox*-cluster evolution in this fish lineage.

**Figure 1.**
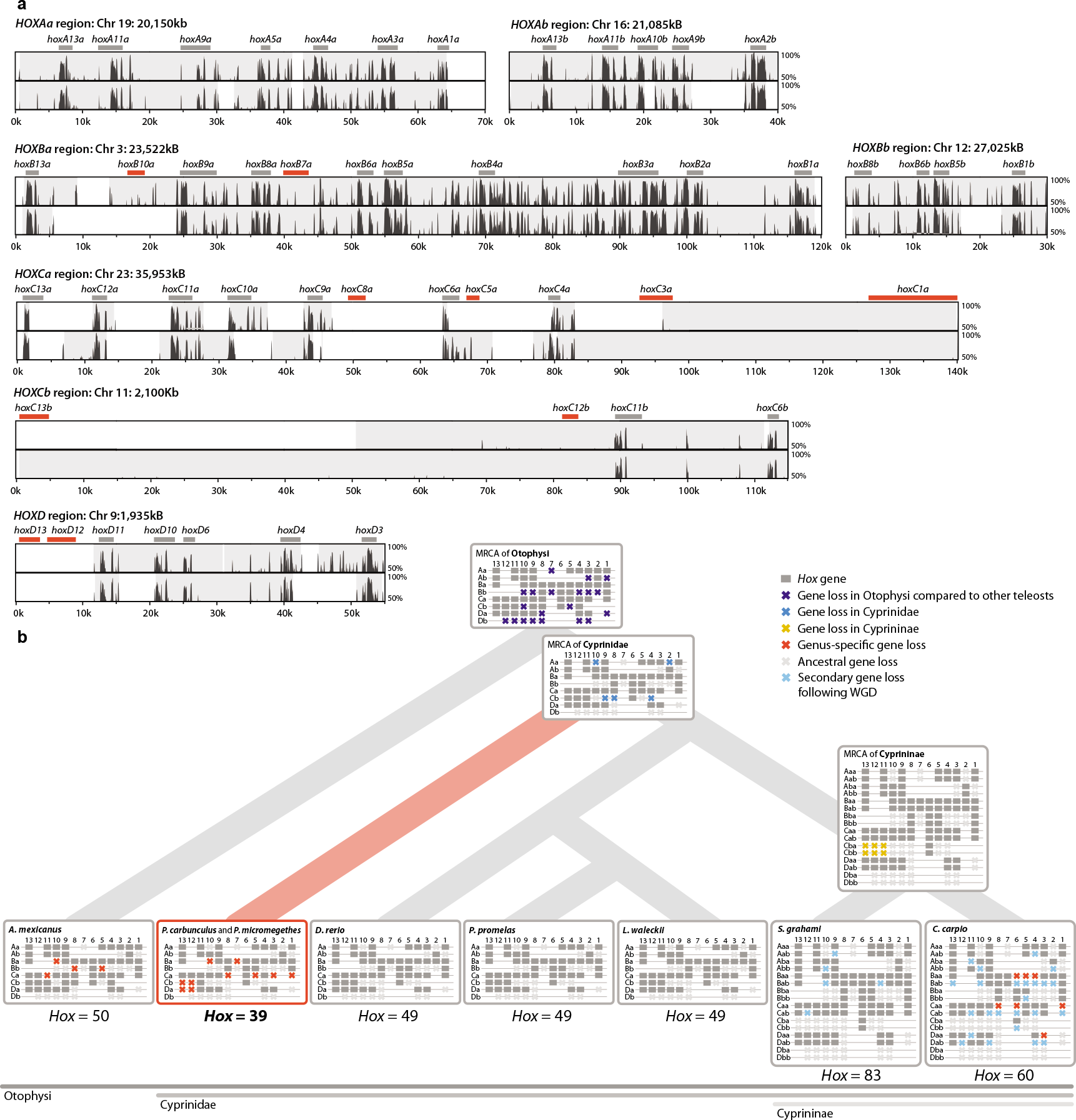
Sequence similarity plots and reconstructed evolutionary history of teleost *Hox* genes. a) Pairwise sequence similarity between zebrafish (*D. rerio*) and *P. carbunculus* (top) and *P. micromegethes* (bottom) for syntenic *Hox* regions. Sequence similarity from 50–100% is shown with green bars representing the location of complete copies of recovered *Hox* genes, while red bars represent the expected location of the unrecovered genes. Grey shading represent contiguous sequences in either of the *Paedocypris* species with sufficient similarity to be mapped to the zebrafish chromosomes. b) Cladogram of selected otophysan species including hypothetical ancestors and ancestral losses (topology according to Stout *et al*. 2017). Genus-specific *Hox* gene losses are shown in red. Only intact gene copies are shown. *Hox* cluster information from other species, and the ancestral states, are based on Henkel *et al*. (2012) ^29^, Pascual-Anaya *et al*. (2013)^30^, Yang *et al*. (2016)^31^, and Ensembl^32^.

Given its progenetic phenotype and the fact that ~20% of the zebrafish *Hox* gene repertoire is apparently absent in *Paedocypris*, we also investigated the presence of other key genes involved in various developmental pathways. We first determined the overlap in gene space between the two *Paedocypris* species, the zebrafish, and the two tetraodontid models *Di. nigroviridis* and *T. rubripes*, as these pufferfishes have similarly small genomes as the *Paedocypris* species. Figure 2a shows the number of shared orthogroups for all species, identified using OrthoFinder^33^, and highlights the number of orthogroups without orthologs in *Paedocypris* (numbers in bold). We used the set of genes from these 1,160 orthogroups to identify a comprehensive list of 1,581 genes that had gene ontologies associated with different system development pathways and pattern specific processes in *D. rerio* (Methods and Supplementary Table 2). These 1,581 genes were then used as queries to screen the genomic sequences and annotated protein sets of *Paedocypris*. Fourteen of these genes, primarily involved in skeletal- (*ucmab, disc1* and *Ibh*), muscle- (*ccdc78* and *marcksa*), and nervous system (*nkx2.9, pho, cntf, thfrsf1b, tnfrsf19* and *lepa*) development, were not found in either of the *Paedocypris* species (Figure 2b). Adjacent flanking genes in zebrafish could, however, be identified in *Paedocypris* for all 14 genes (Supplementary Table 3).

**Figure 2.**
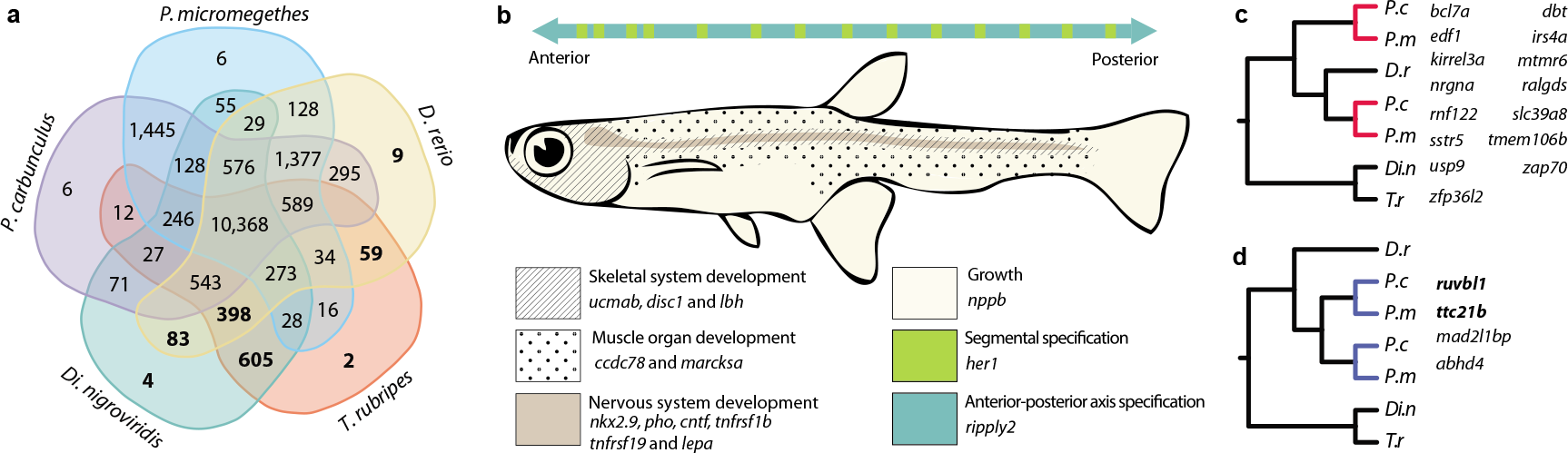
Gene loss and duplication of developmental process genes. a) Five-species comparison of shared orthogroups for the identification of genes lost in *Paedocypris*. Bold numbers represent orthogroups without any *Paedocypris* orthologs. Orthogroups were identified using OrthoFinder^33^ on the basis of full protein datasets for all included species. b) Lost developmental pathway genes and schematic representation of phenotypically affected body segments in *Paedocypris*. c) Gene duplicates retained in *Paedocypris*. d) Genus specific gene duplications. Genes in bold are associated with a truncated phenotype in *D. rerio*.

Because phenotypic changes can also result from gene duplication^34,35^, we identified genes with two copies in *Paedocypris*, but only a single copy in zebrafish and pufferfishes. Based on the topology of gene trees generated for these genes, we differentiated between genes originating from duplication events predating the divergence of *Paedocypris* and *Danio*, where only one copy is retained in zebrafish (Figure 2c), and *Paedocypris*-specific gene duplication events (Figure 2d). Interestingly, although only four genes could be identified as recent duplicates in *Paedocypris*, two of these (*ttc21b* and *ruvbl1*) are associated with deformed phenotypes in zebrafish, including shortening of the anterior-posterior axis and decreased head size^32^.

### Genome miniaturization through intron shortening

In parallel to their miniaturized body size, the two *Paedocypris* species show a surprising evolutionary trajectory in terms of genome miniaturization. Compared to the genome sizes of zebrafish (~1,5 Gb)^7,32^ and other cyprinid fishes (0.81–3.5 Gb)^36^, we find that the two *Paedocypris* species have substantially smaller genomes (0.45–0.52 Gb), yet similar numbers of genes (Table 1). Comparative analyses of vertebrate genomes have shown that genome size reduction is typically characterized by shorter introns and reduced repeat content^37,38^. However, substantial loss of protein-coding genes^38,39^, large segmental deletions facilitated by fission of macrochromosomes^38^, and a reduced rate of large insertions have also been demonstrated to play a role in reducing or constricting genome size^37,40^.

**Table 1.**
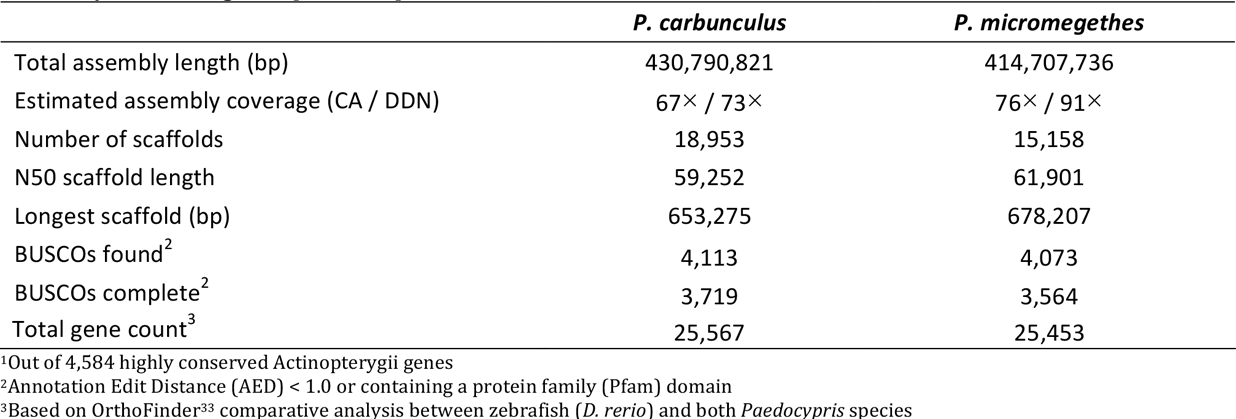
Assembly statistics, gene-space completeness-, and annotation metrics.

As the *Paedocypris* genome sizes are comparable to those of tetraodontid pufferfishes, we compared the gene repertoire of *Di. nigroviridis* and *T. rubripes* to that of *Paedocypris* and *D. rerio* to determine what genomic features are shared among these lineages. In order to obtain a consistent gene set for comparative analysis of gene-, exon-, and intron lengths, we categorized the full gene set of all five species into orthologous groups using the software OrthoFinder^33^. We identified 10,368 orthogroups containing genes from all five species (Figure 2a), which included 16,142 genes in zebrafish, 15,287 in *P. carbunculus*, 14,181 in *P. micromegethes*, 14,529 in *Di. nigroviridis*, and 14,393 in *T. rubripes*. The total gene length and the proportions of exonic and intronic regions for this gene set are shown in Figure 3a. Although the accumulative length of this gene set is significantly smaller in both *Paedocypris* species and the two tetraodontids than in the zebrafish, the proportional change is more subtle, with the gene set constituting on average 29–30% of the total genome length in *Paedocypris* and the tetraodontids and 40% in zebrafish. Further, even though we detected minor differences in this gene set regarding the average number of exons per gene in *Paedocypris* (10.04) compared to *D. rerio* (11.21), the observed 47% reduction in total exon length of *Paedocypris* in relation to zebrafish cannot be attributed to exon loss alone, as the two tetraodontids show very similar results to *Paedocypris* with even higher average exon count (11.89). However, the majority of the total gene length reduction observed in *Paedocypris* and tetraodontids is due to an 80–84% reduction in overall intron size (Figure 2a). To rule out that the overall reduction in intron size is driven by a highly deviant fraction of the *Paedocypris* and tetraodontid gene sets, we also investigated the distribution of gene-, exon-, and intron-lengths of this common gene set (Fig 2b). We observe that zebrafish has substantially fewer short genes compared to all other species, with an average gene length 4.3–5.5 longer than that of *Paedocypris* and tetraodontids. This is not unexpected, as previous studies have reported a lineage-specific expansion of intron size^41^, resulting in an additional peak of intron lengths between 1,000–2,000 bp in zebrafish, shown in Figure 3b. Based on our results, it is apparent that the reduced average gene length in *Paedocypris* is driven by a substantial shift towards consistently shorter introns, similar to, but not as extreme as, in the two tetraodontids. This is further illustrated by the calculated median sizes depicted in Fig. 3b.

**Figure 3.**
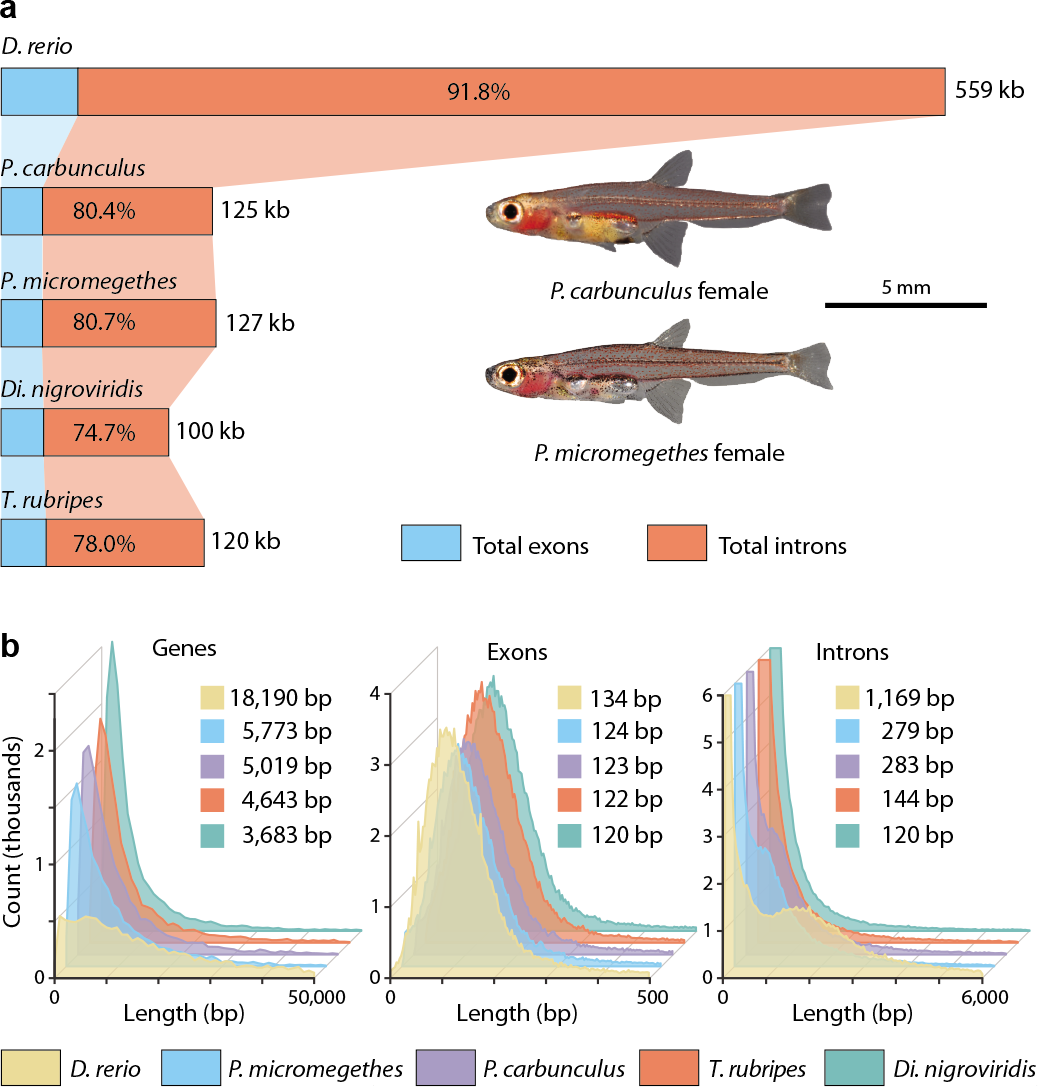
Comparative analyses of repeat content and gene space in zebrafish and *Paedocypris*. a) Total gene length of 14–16,000 genes belonging to common orthogroups, and the proportional contribution of exonic and intronic regions, in zebrafish, *Paedocypris*, and tetraodontids. b) Frequency plot and median values of gene-, exon-, and intron lengths for the common gene set. In all distributions, the values for both *Paedocypris* species are significantly lower than in zebrafish, but significantly greater than those of both tetraodontids. (Wilcoxon Rank Sum test, *p* < 10^−15^). Significant results were not detected for intron- and exon lengths between the two *Paedocypris* species; however, in terms of gene lengths, *P. micromegethes* was reported to be significantly longer.

### Repeat content reduction and genome evolution

The genome size of an organism evolves through the relative rate of insertions and deletions and through the effect of natural selection that either favors or eliminates these changes. Genome shrinkage has thus been postulated to evolve through a bias in the rate of insertions relative to deletions^42^, but the impact of this mechanism, and its importance in genome size evolution is still debated^43^. Importantly, even though deletions indeed appear to be more frequent than insertions, the latter tend to include significantly more base pairs, resulting in the gradual increase in genome size in eukaryotes^42^. Although several other types of mutational activity can promote genome-size expansion, self-replicating mobile elements (*i.e.* transposons) have been identified as the most prominent contributor in this regard^44,45^, and recent studies have confirmed a strong correlation between genome size and the amount of transposable elements (TEs) in teleost fishes and other vertebrates^46,47^. Because the more compact genomes have previously been shown to harbor fewer repeats^38^, we investigated the degree to which this was the case in *Paedocypris*. Using a *Danio-specific* repeat library, we determined the total amount of TEs; DNA-transposons, long- and short interspersed repeats (LINEs and SINEs), and long terminal repeats (LTRs) in the genomes of *Paedocypris*, zebrafish, and four other cyprinid species (*Cyprinus carpio, Pimephales promelas, Sinocyclocheilus grahami*, and *Leuciscus waleckii*) in addition to the characiform cave tetra (*Astyanax mexicanus*) using Repeatmasker^48^. The repeat landscape graphs, illustrating the relative amount of each TE class and the Kimura distance^49^ for each of these are shown in Figure 4a along with a time-calibrated phylogeny of these eight species, as inferred using BEAST^50^ (see Methods). Figure 4a also shows the percentage of each repeat type of various age categories, calculated following Kapusta, *et al*. 2017)^47^. The total number of repetitive elements of each class is shown in Figure 4b, clearly illustrating that even though the proportional differences in repeat content is similar between *Paedocypris* and the other cyprinids, the total number of elements is much lower.

**Figure 4.**
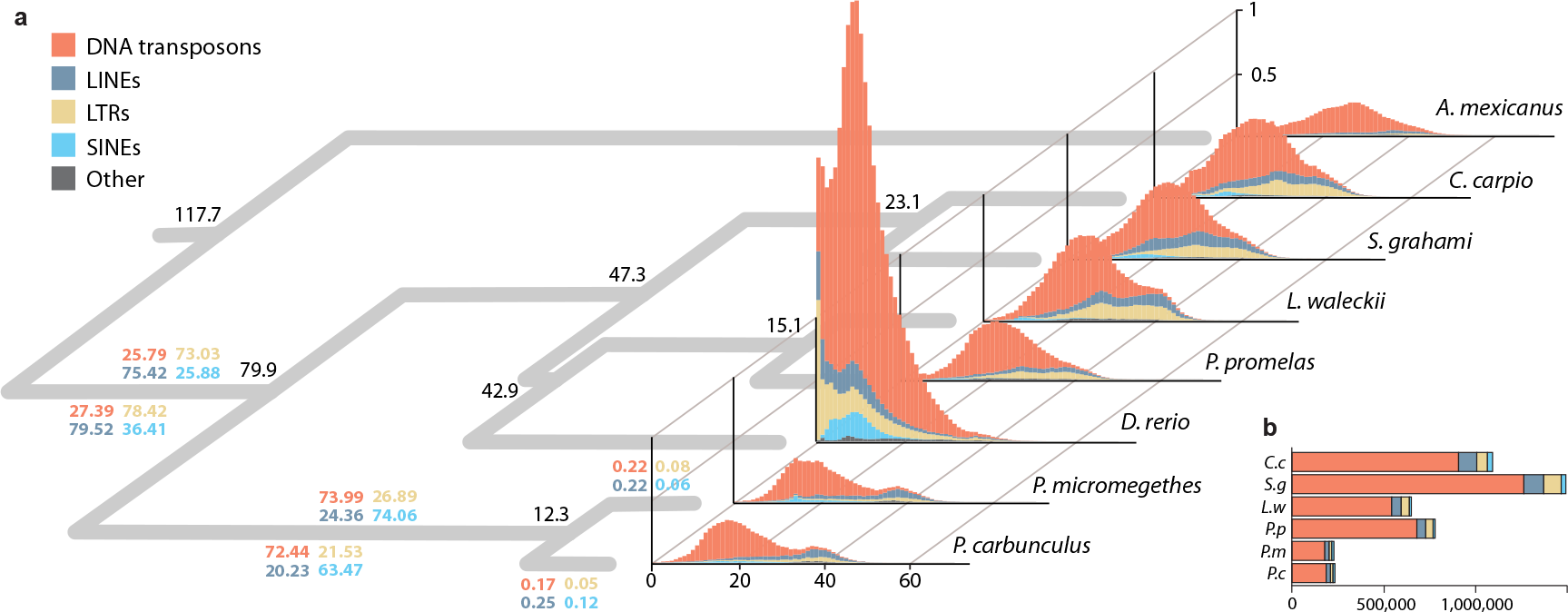
Repeat landscape graphs showing the prevalence and relative age of the four main repeat classes in *Paedocypris* compared to six other teleost genome assemblies. a) Phylogeny with divergence times for the internal nodes for the eight species analyzed. Colored numbers on the branch leading to *Paedocypris* show the percentages of each of the corresponding repeat classes that originate from the time interval corresponding to this branch (117.7–79.9, 79.9–12.3, and more recent than 12.3 Ma). Repeat landscapes represent transposable elements of the four main classes as well as the unclassified ones. The x-axis indicates the Kimura distance^49^ as a proxy for time while the y-axis gives the relative coverage of each repeat class based on the genome size. b) Total number of repetitive elements in each of the cyprinid genomes (excluding zebrafish); colored by class.

Consistent with previous reports^46^, we find the that the zebrafish genome is dominated by repetitive elements, constituting 58.1% of its genome. In contrast, only 15.3 –15.5% of the *Paedocypris* genomes consist of repetitive elements (transposable elements, satellites, simple repeats, and low complexity regions), a percentage that is comparable to that of the other cyprinids investigated (12.3–19.01%). However, in terms of TE content, the *Paedocypris* species have considerably fewer and shorter elements that comprise in total 7.24 and 7.30% of their genomes, compared to the four other cyprinids in which we find on average a genomic proportion of 13.81% (not including zebrafish in which these constitute 53.28% of the genome)(Supplementary Table 4). Based on the timing of the divergence events leading to crown *Paedocypris* and the calculated substitution rate (see Methods), we investigated the proportions of “ancient” and “lineage specific” TEs. Interestingly, only 0.05 – 0.25% of the *Paedocypris* transposons appear to have been incorporated into their genomes after these two species diverged (Figure 4).

As *Paedocypris* thus appears to have reversed the general trend of DNA gain^43,43^ towards gradual DNA loss, as indicated by their low TE content, we investigated whether this could have been achieved through silencing of transposon activity^40^. We thus explored the genomic content of the PIWI-like genes (*PIWIl1* and *PIWIl2*), which are known for silencing transposons in zebrafish and other vertebrates^51,52^. Interestingly, we identified additional copies of *PIWIl1* in both *Paedocypris* genomes, located in a conserved region ~4 Mb upstream of *PIWIl1* on zebrafish chromosome 8, as shown in Figure 5. One of these gene duplications is shared between the two *Paedocypris* species and therefore appears to have evolved prior to their divergence, while an additional, third copy is found exclusively in *P. carbunculus* (Fig. 5a). These lineage-specific duplications suggest that the gradual genome miniaturization has been, at least in part, attained through increased transposon silencing, leading to a bias of DNA loss over DNA gain. This needs to be tested further, utilizing genomic sequences from other closely related cyprinids like *Danionella* and *Sundadanio*. Probably resulting from the lineage-specific whole genome duplication, a single duplicate of the *PIWIl1* gene was also found in *C. carpio*, but no extra copies could be detected in the other cyprinids; *P. promelas, L. waleckii*, or *S. grahami*. No additional copies of *PIWIl2* were detected in either *Paedocypris* or the other cyprinids.

**Figure 5.**
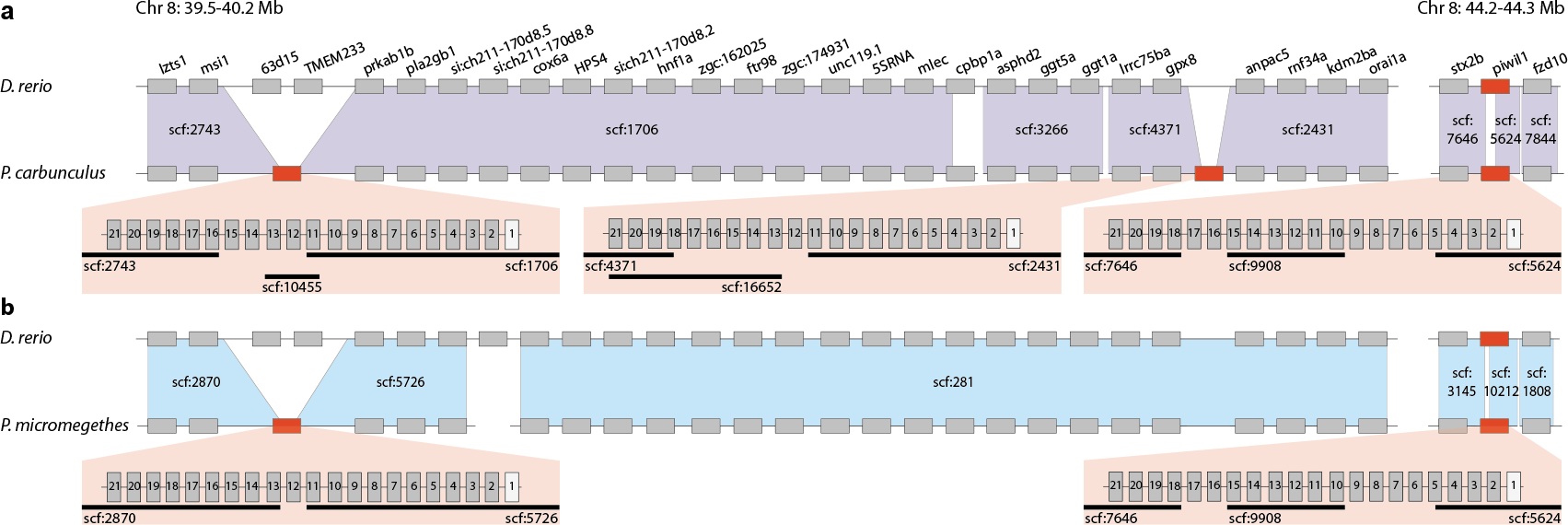
Synteny of the *PIWIl1* loci and surrounding regions in the zebrafish and *Paedocypris* genomes. a) Synteny between zebrafish (top) and *P. carbunculus*, showing the original *PIWIl1* copy (right red rectangle) and the two additional copies (left and middle red rectangles) detected in this genome assembly. b) Same as a) for *P. micromegethes*, showing the single, presumably ancestral, duplicated copy of *PIWIl1* in this genome. Transparent red boxes depict the exon-intron structure of all *PIWIl1* genes in *Paedocypris* with black lines representing the scaffolds covering each region. Not all exons could be recovered for all gene copies.

### Chromosome fusion

Another interesting feature of the miniaturized genome of *Paedocypris* is the reduced number of chromosomes compared to that of zebrafish. While most Cypriniformes have ≥ 24 chromosomes^53^ in the haploid metaphase and zebrafish has 25, *P. carbunculus* only features 15 chromosomes^54^. Since we did not find any indication of large-scale chromosome loss, as would be indicated by substantial gene loss, this discrepancy implies that genome miniaturization in *Paedocypris* spp. has been accompanied by chromosomal fusion since they diverged from their last common ancestor with zebrafish. In order to identify regions on different chromosomes in the zebrafish genome that are now tightly linked in the *Paedocypris* genome, we looked for syntenic regions in *Paedocypris* that contain at least six genes from two different chromosomes in zebrafish. Figure 6 shows the three scaffolds identified in each of the *Paedocypris* species fulfilling these criteria. In both species, nine genes from chromosomes 9 and 21 in zebrafish are co-localized on a single scaffold. For chromosome 11 in zebrafish, we find good evidence in different regions for a fusion with chromosome 1 in *P. carbunculus* but also chromosome 2 in *P. micromegethes*, suggesting that either all three chromosomes may have fused or that species-specific fusion events may have occurred. Although the remaining two putative fusion events can only be detected in one of the species, there are no contradicting results between the two *Paedocypris* species, indicating that fusion of the chromosome pairs 6 and 19 and 22 and 25 is likely to have occurred in the last common ancestor of both species.

**Figure 6.**
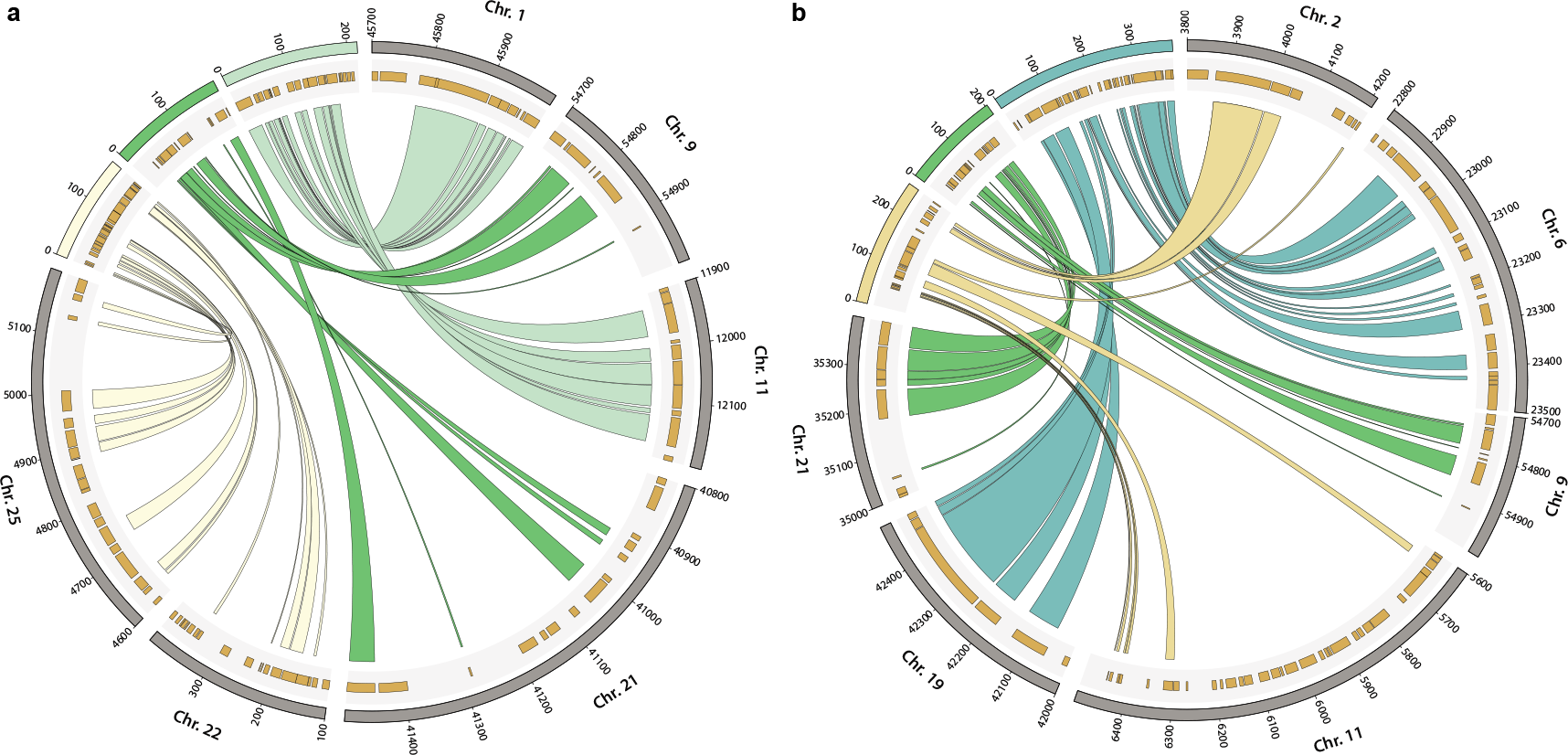
Syntenic regions between zebrafish chromosomes and *Paedocypris* scaffolds. Syntenic regions between zebrafish (*D. rerio*) and *P. carbunculus* (a) or *P. micromegethes* (b). Links between *Paedocypris* scaffolds and D. *rerio* chromosomes indicate orthologous genes. Only the relevant parts of *D. rerio* chromosomes are shown and the locations of all genes in these regions are illustrated in the gray shaded areas. Numbers on scaffolds and chromosomes are given as kbp.

## Discussion

Miniature body size has evolved several times within Cyprinidae^13^. However, little is known about the molecular underpinnings of these miniaturization events, whether these are similar in several instances of miniaturization, and to what extent the parallel evolution of genomic and phenotypic traits is coupled. Given the identification of several classical signatures of genome reduction, including reduced TE content and smaller introns, we conclude that the small *Paedocypris* genomes represent a derived state, having evolved from an ancestor with a substantially larger genome and a larger body size. Reduced genome size has also been reported in other vertebrates^37,38^, yet these reductions include large segmental deletions, chromosome fissions, and massive losses of protein coding genes, which we do not observe in the miniaturized genomes of *Paedocypris*. Moreover, while reduced genome sizes in birds and bats can probably be explained by the adaptation to flight through the need for higher metabolic rates^55^, the causes of genome size reduction in fishes seem in general less apparent. However, the reduced genome sizes in *Paedocypris* may also have evolved as a response to a selection for higher metabolic rate, mediated in this case by its extreme habitat (*i.e.* low oxygen concentrations and pH < 4.0) rather than rapid locomotion.

Small body size has arisen among miniature cyprinids in two different ways, either as proportionate dwarfism, resulting in miniature but otherwise identical copies of their larger ancestors, or through developmental truncation or progenesis leading to larval appearance of the tiny mature fish^13,14^. Evolutionary novelties such as highly modified pelvic and pectoral girdles^14^, an anterior shift in genital pore and anus^13^, and fang-like structures forming from the jaw bones^16^, however, appear to be restricted to developmentally truncated miniatures (*e.g. Paedocypris, Danionella*, and *Sundadanio*). The absence of novelties in proportioned dwarfs would suggest that developmental truncation plays an important role in escaping evolutionary and developmental constraints imposed by the species’ *Bauplan* and in opening up new evolutionary avenues for drastic morphological change^14,16,18^.

This study represents the first report on genomes of highly developmentally truncated fish species. It illustrates how the unique features of *Paedocypris* at the phenotypic level (*i.e.* extreme miniaturization and dramatic developmental truncation) are paralleled at the genomic level, in the form of substantially smaller genomes via reduced intron lengths and lower repeat content, and also through the loss of numerous genes related to development, including highly important *Hox* genes. Although a causal link between miniaturized body size and reduced genome size remains to be established, our study highlights the potential to investigate such connections in a comparative framework, including the sequencing of additional species representing other miniature genera. Nevertheless, these naturally simplified species, having escaped the developmental, and, as we report here, genetic constraints of the cyprinid *Bauplan*, present a novel opportunity to investigate phenotype–genotype relationships in vertebrate development. These naturally occurring extreme phenotypes will greatly aid future research on model organisms and our quest to understand how phenotypic diversity is generated during vertebrate evolution and development.

Recent investigation of the Gulf pipefish genome^56^ has revealed a loss of the last *Hox7* paralog in this species. This has revived the disputed hypothesis that the lack of ribs in both pufferfish and the pipefishes can be attributed to the fact that none of these species have any remaining *Hox7* paralogs^57^. Interestingly, the *Paedocypris* species show a similar loss of the last *Hox7* paralog (*hoxb7a*), yet the two species investigated here do feature ribs. The ribs are, however, reduced and remain poorly ossified^14^, suggesting that although the *Hoxa7/Hoxb7* genes are not the only essential genes for rib development, they do appear to have an influence on their development, as suggested by earlier experiments in mice^58^.

Finally, the observed loss of *Hox13* genes (*Cb* and *Da*) in the two *Paedocypris* species is of special interest as these genes represent the termination of the patterning system. *Hoxd13* was also one of the genes targeted in the successful CRISPR-Cas9 knockout experiment in the zebrafish^59^, that showed that the *Hox* genes play a very similar role in patterning during development of both fins bones, and thus elucidated the evolutionary transition from fins to limbs. We propose that future studies of similar nature should consider using *Paedocypris* species as a complementary system to the zebrafish model, due to its already limited *Hox* gene repertoire, yet relatively close phylogenetic relationship with zebrafish.

## Materials and methods

### Specimens used

*P. carbunculus* was caught at the type locality in Pangkalanbun, Kalimantan Tengah, on the island of Borneo and *P. micromegethes* was caught in Sibu, Sarawak, Malaysia, on the island of Borneo. Both species were caught using dip nets. Immediately upon capture, specimens were killed by an overdose of anesthesia (MS222) following guidelines by the American Society of Ichthyologists and Herpetologists (ASIH) (http://www.asih.org/pubs/; issued 2013). Individuals were preserved in 96% ethanol for subsequent DNA analyses.

### DNA isolation

A single whole specimen of each *Paedocypris* species, stored on 96% ethanol, was used for isolation of high molecular genomic DNA with the EZNA Tissue DNA Kit (Omega Bio-Tek, Norcross, Gerogia, USA), following manufacturer’s instructions. Sample identifiers were LR12004 (*Paedocypris carbunculus*) and LR7898 (*Paedocypris micromegethes*), supplied by Lukas Rüber.

### Sequencing library preparation

Genomic DNA samples were fragmented to lengths of ~550 bp by sonication on a Covaris E220 (Life Technologies, Carlsbad, California, USA) with the following settings: 200 cycles for 45 seconds with Peak Incident Power of 175 W and frequency sweeping mode. All sequencing libraries were constructed following Illumina’s TrueSeq PCR-free library preparation protocol for 550 bp fragments.

### Whole genome sequencing

Based on expected genome sizes of 315-350 Mb^54^, both *Paedocypris* genomes were sequenced to ~90× coverage on the Illumina HiSeq 2500 platform, with the Illumina 500 cycles kit (Rapid mode) with on-board clustering. Per species, this produced 151-155 M paired reads of 250 bp each.

The Kapa Library quantification kit for Illumina (Kapa Biosciences, Wilmington, Massachusetts, USA) was used to find the correct molarity (nM) before sequencing.

### Genome assembly

Both *Paedocypris* genomes were initially assembled using two different assembly programs, the “Overlap-Layout-Consensus” based Celera Assembler^60^ and the “de Bruijn graph” based DISCOVAR *de novo*^61^. We then used the Metassembler software^62^ to merge and optimize these two assemblies, producing a reconsolidated single assembly with superior quality for each of the two *Paedocypris* species (Table 1, Supplementary Note, and Supplementary Table 5).

### Assembly quality assessment

The quality of the different genome assemblies was assessed by comparing the proportion of conserved genes detected, as a measure of gene-space completeness. We used the program BUSCO v2.0 (Benchmarking Universal Single-Copy Orthologs)^63^, which searches for 4,584 highly conserved single-copy actinopterygian genes. Results are listed in Table 1.

We also assessed the assembly quality of the three different assembly versions with the software FRC^bam^ [64], which identifies “features” (incorrectly mapped reads, incorrect insert size, coverage issues, etc.) in each of the assemblies, and ranks the different assembly versions according to the number of detected features. Additional information on the execution of these programs is available in the Supplementary Note and the resulting graphs are presented in Supplementary Figs. 1 and 2.

### Annotation

Structural and functional annotation of both reconsolidated *Paedocypris* genomes was performed with two iterative rounds of the MAKER2 (v 2.31.8)^65–67^ pipeline, following the instructions by Sujai Kumar (available at https://github.com/sujaikumar/assemblage/blob/master/README-annotation.md). Numbers of genes annotated are listed in Table 1. See Supplementary Note for in-depth descriptions of additional software and commands used.

### *Hox* gene search

In addition to the two *Paedocypris* genomes, *Hox* gene content was also investigated in the genomes of *Pimephales promelas* (GCA_000700825.1_FHM_SOAPdenovo_genomic.fna) and *Leuciscus waleckii* (GCA_900092035.1_Amur_ide_genome_genomic.fa) based on BLAST^68^ searches, using 70 zebrafish *Hox* transcripts from Ensembl as queries (including some truncated variants and isoforms), representing the 49 unique *Hox* genes (Supplementary Table 1). The similarity stringency threshold used in these searches was 1e^−20^. We also conducted additional searches using Exonerate v2.2.0^69^ for genes whose presence could not be established based on the BLAST search. For those genes that could not be detected using either method, additional searches were conducted using the orthologous protein sequence from the cave tetra (*Astyanax mexicanus*). Contiguous sequences from both *Paedocypris* genomes were extracted from the scaffolds assembled with Metassembler, spanning all hit regions plus 10 kb upstream and downstream of each hit. These scaffold sequences were aligned to the orthologous sequences extracted from the zebrafish genome (GRCz10) using mVista^70^.

### Identification of lost and expanded developmental genes

In order to obtain a complete list of genes from *D. rerio* that were associated with various developmental pathways that could be compromised in *Paedocypris*, we started out with the orthogroups found not to contain orthologs from either *Paedocypris* species, as identified using OrthoFinder^33^. We further utilized the gene ontology information associated with all genes belonging to these orthogroups, and identified key gene ontology terms: GO:0009948 (anterior–posterior axis formation), GO:0009950 (dorsal–ventral axis formation), GO:0040007 (growth), GO:0007517 (muscle organ development), GO:0007399 (nervous system development), GO:0001501 (skeletal system development), and GO:0007379 (segment specification). This data set contained 1,581 unique genes from *D. rerio*, and 64 of these genes could not be detected in the *Paedocypris* genome sequences using TBLASTN^68^ with a similarity cutoff of 1e^−20^, and their presence was further examined by running Exonerate^69^ (v2.2.0). Based on the Exonerate results, reciprocal BLAST searches, annotation, and identification of flanking genes, we could confidently determine that 14 of these genes were indeed not present in either of the *Paedocypris* genomes. Using expression data from Ensembl^32^, we reconstructed the hypothetically affected body segments in *Paedocypris*, as illustrated in Figure 2b.

We further investigated whether we could confidently identify developmental genes that are now duplicated in *Paedocypris* but not in *D. rerio.* From the full list of orthogroups, we identified 138 groups that were represented by a single copy in the two tetraodontid pufferfishes and in *D. rerio*, but by two apparent copies in both *Paedocypris* species. The alignments of these genes were then screened for missing data, and based on manual inspection of these alignments, only alignments with < 35% missing data were included in further analysis. Out of the 35 genes that met this criterion, 19 could be confirmed through annotation, and had a gene tree consistent with the two hypotheses outlined in Figure 2c and 2d.

### Gene space evolution

In order to assess the changes of gene-, exon-, and intron sizes in *Paedocypris* we first identified the proteomic overlap of *Paedocypris, D. rerio, Di. nigroviridis*, and *T. rubripes* by running the software OrthoFinder^33^ on the complete protein sets of these five species. We used the full protein sets from Ensembl^32^ (v. 80): *D. rerio* (GRCz10), *Di. nigroviridis* (TETRAODON8), and *T*. rubripes(FUGU4). However, as some of the *D. rerio* genes have more than one protein or transcript in the Ensembl database, the output from BioMart (31,953 genes and 57,349 proteins) was filtered so that only the longest protein sequence from each gene was used in the analysis, and genes without protein sequences were removed. This resulted in a set of 25,460 genes with a single protein prediction. For the *Paedocypris* species, the “standard” gene sets resulting from the annotation were used as input. These sets were filtered to include only genes with AED (Annotation Edit Distance) scores < 1 or those with a Pfam domain. By using only genes belonging to the 10,368 orthogroups found to contain orthologs from all these species (Fig. 2a), we obtained a comprehensive but conservative dataset as the basis for these analyses. Information about each of the corresponding genes in *D. rerio, Di. nigroviridis*, and *T. rubripes* was obtained from BioMart, and included the Ensembl gene and protein ID, and the chromosome name in addition to the start and stop position for each gene, transcript, and exon. Intron sizes were then calculated on the basis of exon positions, using a custom script (“gene_stats_from_BioMart.rb”). In some cases, the sum of exons and introns did not equal the total length of a gene, which appears to be caused by inconsistency in the registration of UTR regions in the Ensembl database for individual genes. In these cases, to be conservative with regard to intron length estimates, the gene length was shortened to correspond to the sum of the exons and the corresponding introns between these.

Intron and exon lengths for the two *Paedocypris* species were calculated in a similar manner, but on the basis of the “standard” filtered annotation file in “gff” format produced as part of the annotation pipeline. Also for these species, the intron lengths were determined on the basis of identified exons, with another custom script (“gene_stats_from_gff.rb”). Gene-, exon-, and intron length histograms were plotted with the R package ggplot2. Custom Ruby scripts are available for download at https://github.com/uio-cees/Paedocypris_gene_stats

### Repeat content analysis

The repeat contents of the two *Paedocypris* genomes and the model organism genomes were assessed using RepeatMasker (v4.0.6)^48^ with the Danio-specific repeat library included in the program, and with the “-s” setting to increase sensitivity. The following model organism genome assemblies were used in this comparison: *D. rerio* (Danio_rerio.GRCz10.dna.toplevel.fa), *P. promelas* (GCA_000700825.1_FHM_SOAPdenovo_genomic.fna), *L. waleckii* (GCA_900092035.1_Amur_ide_genome_genomic.fa), *Sinocyclocheilus grahami* (GCA_001515645.1_SAMN03320097.WGS_v1.1_genomic.fna), *Cyprinus carpio* (GCA_001270105.1_ASM127010v1_genomic.fna), and *A. mexicanus* (Astyanax_mexicanus.AstMex102.dna.toplevel.fa). Repeat landscape graphs (Fig. 4a) were plotted with the R package ggplot2 based on the “.aligned” output file from RepeatMasker. The proportion of repeats originating from specific time intervals was calculated using the parseRM.pl script^47^ (available at https://github.com/4ureliek/Parsing-RepeatMasker-Outputs), using the *Paedocypris-specific* substitution rate and the “‒‒age” setting set to “12.31,79.98”, according to divergence times estimated with BEAST2. All four repeat classes were analyzed individually using the “‒‒contain” setting.

### Calculation of substitution rate for *Paedocypris*

The *Paedocypris-specific* substitution rate was calculated on the basis of a whole genome alignment of the two *Paedocypris* species. These alignments were created by mapping the sequencing reads of *P. micromegethes* to the *P. carbunculus* assembly using BWA (Burrows-Wheeler Aligner) (v. 0.7.12)^71^ and SAMTOOLS (v. 1.3.1)^72,73^. The number of nucleotide differences in each of the 18,953 alignments (one per scaffold) was identified using a custom script (“find_variable_sites.rb”, available for download at https://github.com/uio-cees/Paedocypris_gene_stats). The substitution rate per million years (0.001288) was then calculated as the sum of total differences (13,415,043) divided by the total number of aligned sites (422,829,036) and two times the estimated crown age of *Paedocypris* (12.31), as inferred by the BEAST v.2.4.5^50^ analysis.

### Phylogenetic inference of selected otophysan species

To allow the estimation of divergence times between the two *Paedocypris* species and six other otophysan taxa for which genomic resources were available (*D. rerio, A. mexicanus, C. carpio, P. promelas, S. grahami*, and *L. waleckii*), we followed the pipeline for phylogenetic marker selection presented in Malmstrøm et al. (2016 and 2017)^74,75^ with few modifications. These modifications included the following changes: Marker selection began with a set of 3,238 cave fish (A. *mexicanus*) exons, for which at least five orthologs were known among the seven species *D. rerio, Gadus morhua, Gasterosteus aculeatus, Oreochromis niloticus, Oryzias latipes, Poecilia formosa*, and *T. rubripes*, according to version 87 of the Ensembl database. This marker set was then used to identify potential orthologs from the genomes of the eight selected otophysan taxa based on TBLASTN^68^ searches followed by a strict filtering procedure. Compared to Malmstrøm et al. (2016), we applied a lower dN/dS threshold of 0.25 to exclude markers potentially affected by positive selection, and we removed all markers for which no homologs could be detected in one or more of the eight otophysan genomes. We also applied stricter thresholds on clock-like evolution of candidate markers, so that all genes with an estimated coefficient of rate variation above 0.8 as well as those with a mean mutation rate above 0.0004 per site per million years were excluded. We identified 138 genes with a total alignment length of 135,286 bp which were subsequently used for analysis with BEAST v.2.4.5^50^. Since the topology of otophysan taxa has previously been resolved with a more comprehensive phylogenetic dataset^21^, we here focused on the inference of divergence times only, by using the topology inferred by Stout et al. (2016) as a starting tree and excluding all of BEAST2’s operators on the tree topology. Divergence times were estimated by calibrating the most recent common ancestor of Cypriniformes and Characiformes with a lognormal distribution centered at 121 Ma (standard deviation on log scale: 0.1) according to the results of Malmstrøm et al. (2016). We performed two replicate BEAST2 analyses with 800 million MCMC iterations, of which the first 100 million were discarded as burnin. Convergence was assessed based on similarity of parameter traces between run replicates and effective sample sizes (ESS) greater than 200. A maximum clade credibility (MCC) summary tree with node heights according to mean age estimates was produced with TreeAnnotator v.2.1.2^50^.

### Identification of PIWI-like genes

We investigated the presence of PIWI-like genes in the genomes of the two *Paedocypris* species and the other cyprinids using Exonerate^69^ (v2.2.0) with the longest transcripts available for the two PIWI-like homologs from zebrafish; *PIWIl1* (ENSDARG00000041699) and *PIWIl2* (ENSDARG00000062601). Regions containing sequences spanning more than three introns were aligned to the zebrafish exons using mafft^76^ as implemented in AliView^77^ (v1.17.1). Intron sequences were aligned manually based on the established exon structure, using the full-length scaffold sequences. Local gene synteny to zebrafish chromosome 8, surrounding the putative *PIWIl1* copies, was confirmed through reciprocal BLAST searches using both the MAKER2 annotated proteins and proteins predicted by GeneScan (online version)^78^ as queries.

### Identification of chromosomal rearrangements (fusions)

As *P. carbunculus* has been shown to have a haploid chromosome count of 15 [^54^], potential chromosomal fusions were investigated on the basis of disrupted synteny of zebrafish chromosomes in relation to *Paedocypris*.

We identified putative homologous regions between the zebrafish genome assembly and each of the *Paedocypris* species’ genome assemblies by using MCScanX^79^. In short, the predicted proteins for each *Paedocypris* species were merged with predicted proteins from zebrafish into a single file, and BLASTP^68^ was executed with this file as both query and target, thus identifying putative homologs both within each species and between. The genomics positions of the proteins were extracted from the annotation files, and the BLASTP results and the genomic positions were provided to MCScanX for identifying the putative homologous regions.

## Acknowledgments

Fieldwork in the peat swamp forests in Malaysia and Indonesia was funded by the Natural Environmental Research Council (NERC; NE/F003749/1, to L.R. and R.B.), National Geographic (8509-08, to L.R.), and the North of England Zoological Society (to L.R.). Fieldwork in Sarawak was conducted under permits issued by the Economic Planning Unit, Prime Minister’s Department, Malaysia UPE 40/200/19/2534) and the Forest Department Sarawak (NCCD.970.4.4[V]-43) and fieldwork in Indonesia was conducted under permits issued by the Indonesian Institute of Sciences (LIPI) and the Kementerian Negara Riset dan Teknology (RISTEK; 3/TKPIPA/FRP/SM/III/2012). We thank E. Adamson H. Budianto H. Ganatpathy, S. Lavoué, M. Lo, H. Michael, and S. Sauri for their help in the field. All computational work was performed on the Abel Supercomputing Cluster (Norwegian metacenter for High Performance Computing (NOTUR) and the University of Oslo) operated by the Research Computing Services group at USIT, the University of Oslo IT-department. Sequencing library creation and high-throughput sequencing were carried out at the Norwegian Sequencing Centre (NSC), University of Oslo, Norway. This work was funded by grants from the Naturhistorisches Museum der Burgergemeinde Bern to L.R. and the Research Council of Norway (RCN grants 199806 and 222378) to K.S.J. H.H.T acknowledges funding from the National University of Singapore (NUS, R-154-000-318-112) and Lee Kong Chian Natural History Museum. W.S. acknowledges funding from the European Research Council (ERC) and the Swiss National Science Foundation (SNF).

